# Subunit vaccination enhances protection conferred by prior *Mycobacterium tuberculosis* exposure

**DOI:** 10.64898/2026.07.10.737547

**Authors:** Sara B. Cohen, Joshua S. Woodworth, Thomas Lindenstrøm, Anele Gela, Fergal J. Duffy, John D. Aitchison, Thomas J. Scriba, Elisa Nemes, Rasmus Mortensen, Kevin B. Urdahl

## Abstract

*Mycobacterium tuberculosis* (Mtb) remains the leading cause of death from a single infectious agent. Although BCG protects against disseminated tuberculosis in infants, it has failed to curb transmission among adults. Animal models are used to prioritize vaccines entering clinical trials, but current models use Mtb-naïve mice, limiting their relevance to humans in endemic settings, the primary targets of vaccine trials. Here we used a mouse model of Mtb exposure, CoMtb, where a cervical lymph node Mtb infection protects against aerosol challenge, consistent with historical observations in humans that prior Mtb exposure reduces disease risk. When we compared the impact of CoMtb on vaccine efficacy, BCG failed to provide added protection over CoMtb, while protein subunit vaccination further reduced lung burdens. This protection was associated with decreased KLRG1+ vascular CD4+ T cells and enhanced polyfunctional lung CD4+ T cells before and after aerosol challenge. Our findings were corroborated in an analysis of vaccinated humans stratified by QuantiFERON-TB status, which revealed a similar enrichment of polyfunctional CD4+ T cells in Mtb-exposed individuals. These data support the use of CoMtb to test vaccine efficacy in Mtb-exposed individuals and highlight the potential of subunit vaccines in endemic regions.

## Introduction

Tuberculosis remains a leading cause of infectious morbidity and mortality despite widespread BCG vaccination. The efficacy of BCG has been strikingly variable, ranging from 0-80% depending on the population, and this variation has been attributed to several factors, including age, latitude, and prior mycobacterial exposure, among others(1, 2). While BCG is protective in infants, especially against disseminated disease, it has been ineffective in curtailing TB disease in adolescents and adults. Because Mtb is transmitted by adolescents and adults with TB disease, but negligibly by infants and young children(3), BCG has been ineffective at stopping Mtb transmission and curbing the TB pandemic. As a result, a large proportion of those living in high transmission settings have been exposed to Mtb by adolescence. Because of the desire to develop a vaccination regimen that curbs Mtb transmission, most current vaccine trials target adolescents and young adults, the primary Mtb transmitters. This creates a confounding scenario, as candidate vaccines are prioritized for clinical trials based on pre-clinical vaccine testing in Mtb-naïve animals, while many individuals in vaccine-targeted human populations have pre-existing immunity due to prior Mtb exposure. Because Mtb-naïve animals likely have very different immune landscapes than those that are Mtb-sensitized, new animal models are needed to test TB vaccine efficacy in the context of pre-existing immunity.

Most animal models for testing vaccines in the setting of prior Mtb exposure rely on an initial aerosol Mtb challenge (∼50 CFU) followed by 6-12 weeks of antibiotic treatment and a subsequent rest period prior to vaccination(4–7). This contrasts with Mtb re-infection in asymptomatic humans previously exposed to Mtb because the mice experience a 6-week period of “active” lung disease that could dramatically influence lung immune responses. There is growing evidence that re-infection represents the most common pathway to active TB in endemic regions(8); thus, a platform that could assess a vaccine’s ability to confer protection to new infection in the context of pre-existing immunity is highly desirable. To address this gap, we have taken advantage of a recently described model in which mice are infected in the dermis of the ear with Mtb, which induces a chronic infection of the cervical-draining lymph node, subsequently termed the concomitant Mtb (CoMtb) model(9–11). Mice in this model rarely have detectable bacteria in the lung yet retain live bacteria in the lymph node for at least one year(10). This reflects findings in post-mortem studies from the pre-antibiotic era that lymph nodes are a common site for Mtb persistence in asymptomatic individuals even when Mtb cannot be detected in the lungs(12). This model has been previously used to study factors associated with TB “reactivation”(9, 13), concomitant immunity against subsequent aerosol challenge(10), and remodeling of myeloid cells(11, 14), but it has not previously been used as a platform for vaccine testing. A critical question that remains in the field is what properties of a vaccine could improve upon the protection conferred by chronic Mtb lymph node infection against subsequent aerosol challenge.

In this study, we performed immunogenicity and protection studies using two protein subunit vaccines, H56 and H107, in the setting of pre-existing immunity with the CoMtb mouse model. These two vaccines each express Mtb protein antigens that, in this study, are delivered within a cationic liposomal adjuvant, CAF01, a member of the CAF family of adjuvants that expresses dimethyldioctadecylammonium (DDA) and the Mincle agonist trehalose-6,6’-dibehenate (TDB) to promote a mixed (Th1/Th17) and long-lasting T cell response in human PBMC(15–17) as well as in mice(16, 18–20). Another CAF member, CAF10b, expresses both TLR9 and Mincle agonists to induce long-lasting Th1 and Th17 responses in mice and NHP that are superior to the responses induced by CAF01 in head-to-head experiments (21). For this reason, CAF10b has recently progressed to a Phase I clinical trial in humans using the H107 vaccine. Human trials completed thus far using these subunit vaccines, including the H56 trial reported in this study, have used the IC31 adjuvant(15, 22–27), which activates the TLR9/MyD88 pathway to induce a strong Th1 response(28). Here we find that each subunit vaccine performed well in both Mtb-naïve and CoMtb post-exposure models of murine Mtb infection, whereas BCG was only effective in the Mtb-naïve setting. The protection conferred by subunit vaccination in the post-exposure setting was associated with enhanced CD4 T cell polyfunctionality, particularly expression of IL-17, and reduced terminal differentiation. These protective features were present in the spleens and lungs of Mtb-exposed mice even prior to aerosol challenge, suggesting that subunit vaccination, but not BCG, skews the peripheral immune landscape toward protective responses that are remarkably enhanced rather than impaired by prior Mtb infection. Finally, our findings of enhanced immune features in the murine CoMtb model were validated in a clinical cohort of QuantiFERON-TB (QFT)+ or QFT-humans with or without H56 immunization with the IC31 adjuvant. Together, these data illustrate the promise of protein subunit vaccines for use in adult populations who may have already experienced Mtb infection and underscore the utility of the CoMtb model of pre-existing immunity as a preclinical testing platform for vaccine candidates.

## Results

### BCG efficacy is restricted to mycobacteria-naïve mice

There is longstanding speculation in the field that prior mycobacterial exposure is a major driver of poor BCG efficacy(1). To experimentally address whether prior Mtb exposure influences BCG-mediated protection, we took advantage of the CoMtb mouse model, wherein a chronic infection is established in the cervical draining lymph node with minimal to no dissemination of bacteria to the lung, thus mimicking Mtb exposure in the absence of active pulmonary disease(9, 10). In these experiments, control C57BL/6xBalbc F1 (CB6F1) mice were either intradermally infected (CoMtb) or BCG-immunized, and a third group was administered CoMtb 6 weeks prior to BCG to model immunization in the setting of ongoing Mtb infection. Eight weeks after the final immunization, mice were aerosol challenged with H37Rv Mtb, and lungs were harvested 4 weeks post-infection (Fig 1A). As expected, BCG immunization provided ∼1 log of protection in the lung compared to unvaccinated mice, and as previously described, CoMtb was similarly protective at this time point, with a slight increase in protection relative to BCG(10). Interestingly, BCG in the setting of ongoing Mtb exposure failed to provide any additional protection over that already observed by CoMtb alone (Fig 1B). This is consistent with the observation in humans that BCG is effective in Mtb-naïve individuals and infants but fails to provide protection amongst those living in endemic areas or amongst individuals who are tuberculin skin test (TST)+/QFT+(1, 2).

**Figure 1.**
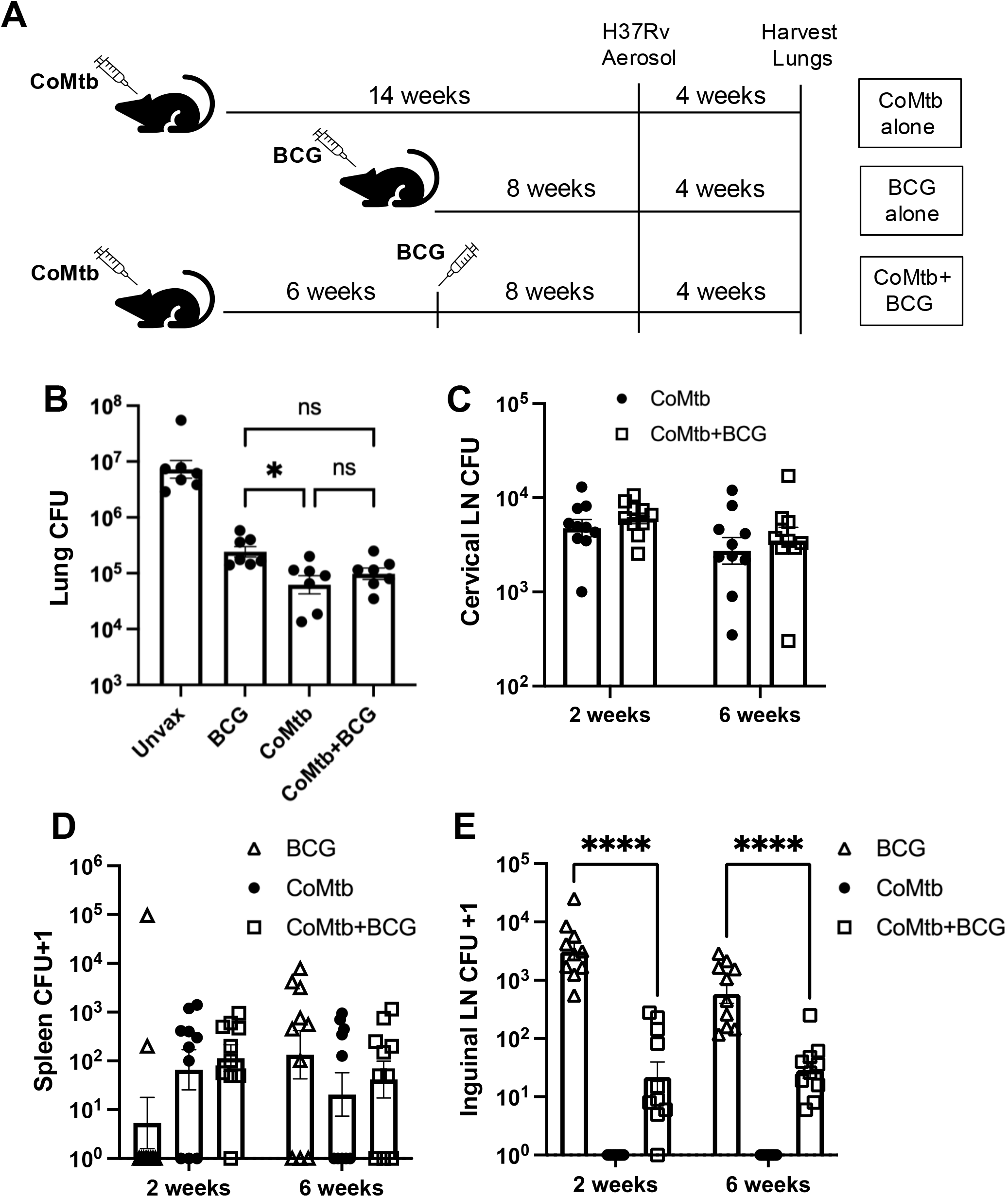
BCG-mediated protection is restricted to naïve individuals and is abrogated in the context of ongoing Mtb infection. (A) Schematic of the experimental set-up. Briefly, the contained Mtb (CoMtb) model was initiated by intradermal injection of 10^4^ virulent H37Rv in the ear of CB6F1 mice. Six weeks later, mice were either left unvaccinated or immunized with 10^6^ BCG subcutaneously. Eight weeks following BCG, mice were aerosol-challenged with H37Rv. (B) Left lung lobes were harvested for CFU analysis 4 weeks post-aerosol challenge. (C-E) In some experiments, the cervical draining lymph node (cLN) (C), spleen (D), and inguinal LN (iLN) (E) were harvested after CoMtb alone, BCG alone, or CoMtb followed by 2 and 6 weeks of BCG, all prior to aerosol challenge. CFU values do not distinguish between Mtb and BCG. Log-transformed CFU data are shown as the means ± SEM. Statistical analysis was performed by one-way ANOVA (B) or two-way ANOVA (E); *, p<0.05, ****, p<0.0001.

We next asked whether the failure of BCG to protect previously exposed mice in this model was due to either the masking or blocking hypothesis, two schools of thought that are commonly used to explain the lack of BCG efficacy in humans(29). In the former, a pre-existing immune response to mycobacteria prevents noticeable additive protection from any potential beneficial effects of BCG, whereas the latter predicts that a prior mycobacterial infection prevents the establishment of BCG within the host. To test this, CB6F1 mice were administered CoMtb or BCG as controls. In the experimental group, BCG was administered 6 weeks after CoMtb, and inguinal lymph nodes (iLN, which drain the BCG immunization site), spleen, and cervical draining LN (cLN, which drain the intradermal CoMtb inoculation site) were harvested at 2 and 6 weeks post-BCG immunization. The presence of BCG had no impact on the levels of CoMtb-derived Mtb found in the cLN at either timepoint (Fig 1C). BCG was rarely found in the spleen at 2 weeks (2 of 10 mice), and the levels of Mtb derived from the intradermal infection were unaffected by the presence of BCG. More BCG was recovered in the spleens of the BCG-only control group after 6 weeks, but the presence of concomitant CoMtb did not significantly change the overall splenic bacterial burdens; although we did not distinguish between BCG and Mtb colonies in this experiment, there was likely a mix of the two bacterial strains at the later timepoint (Fig 1D). However, whereas all BCG-immunized mice had ∼10^3^ CFU in the iLN following BCG vaccination, this bacterial load of BCG was reduced by ∼2 logs and ∼1 log after 2 and 6 weeks, respectively, in the presence of CoMtb; importantly, because CoMtb itself was never detected in the iLN of CoMtb-only control mice, the detectable bacteria in the experimental group were likely derived from BCG (Fig 1E). These findings suggest that pre-existing immunity to Mtb prevents the long-term persistence of BCG, providing support for the BCG blocking hypothesis.

### Subunit vaccination provides enhanced efficacy in the setting of prior Mtb infection

For the first time in decades, multiple TB vaccines are currently in various stages of clinical development. Several candidates in the clinical pipeline are protein subunit vaccines, which have been shown to retain efficacy in the context of prior BCG immunization(4) or in Mtb reactivation models(4–6). Given the need for vaccines that can be administered in post-exposure settings, we were interested in whether subunit vaccination might afford greater protection than BCG in the setting of CoMtb. To address this, we used the H56 subunit vaccine, which contains ESAT6, Ag85B, and Rv2660c antigens, in conjunction with the CAF01 liposomal adjuvant, as it has been shown to induce strong polyfunctional T cell responses and long-lasting immunity to subsequent challenge in mice(4, 19). In humans, H56 and a similar vaccine, H1, have been paired with the IC31 adjuvant, showing moderate immunogenicity and strong safety profiles(17, 26, 30). The use of CB6F1 mice in these experiments allowed us to distinguish between vaccine-and infection-driven responses because both ESAT6, which is present in both Mtb and the vaccine, and TB10.4, which is present in Mtb but not the vaccine, contain an MHCII-restricted CD4 T cell epitope recognized by these mice; thus, responses that are unique to ESAT6 stimulation and absent upon TB10.4 stimulation represent vaccine-induced signals.

Given previous data indicating a role for antigen dose in driving effective vs ineffective vaccine responses(5, 7, 23, 25, 31), our initial studies included low (0.05 µg) and high (50 µg) doses of H56 in CAF01 with or without prior Mtb exposure as established by the CoMtb model. As previously reported(5), both high and low doses of H56/CAF01 were protective against aerosol Mtb challenge early following infection in Mtb-naïve mice (Fig 2A). However, at late time points (∼100 days post-infection), only low dose H56/CAF01 was protective, as high dose H56/CAF01 failed to provide any protection compared to unvaccinated mice (Fig 2B). As expected from Nemeth et al.(10) and consistent with Fig 1B, CoMtb was highly protective against aerosol challenge, both at early and late time points (Fig 2A,B). Interestingly, mice that received low dose H56/CAF01 in the setting of CoMtb had significantly lower lung bacterial burdens than mice that received either H56/CAF01 or CoMtb alone at both time points, but high dose H56/CAF01 was ineffective at curbing the lung burden compared to CoMtb at these same time points (Fig 2A,B).

**Figure 2.**
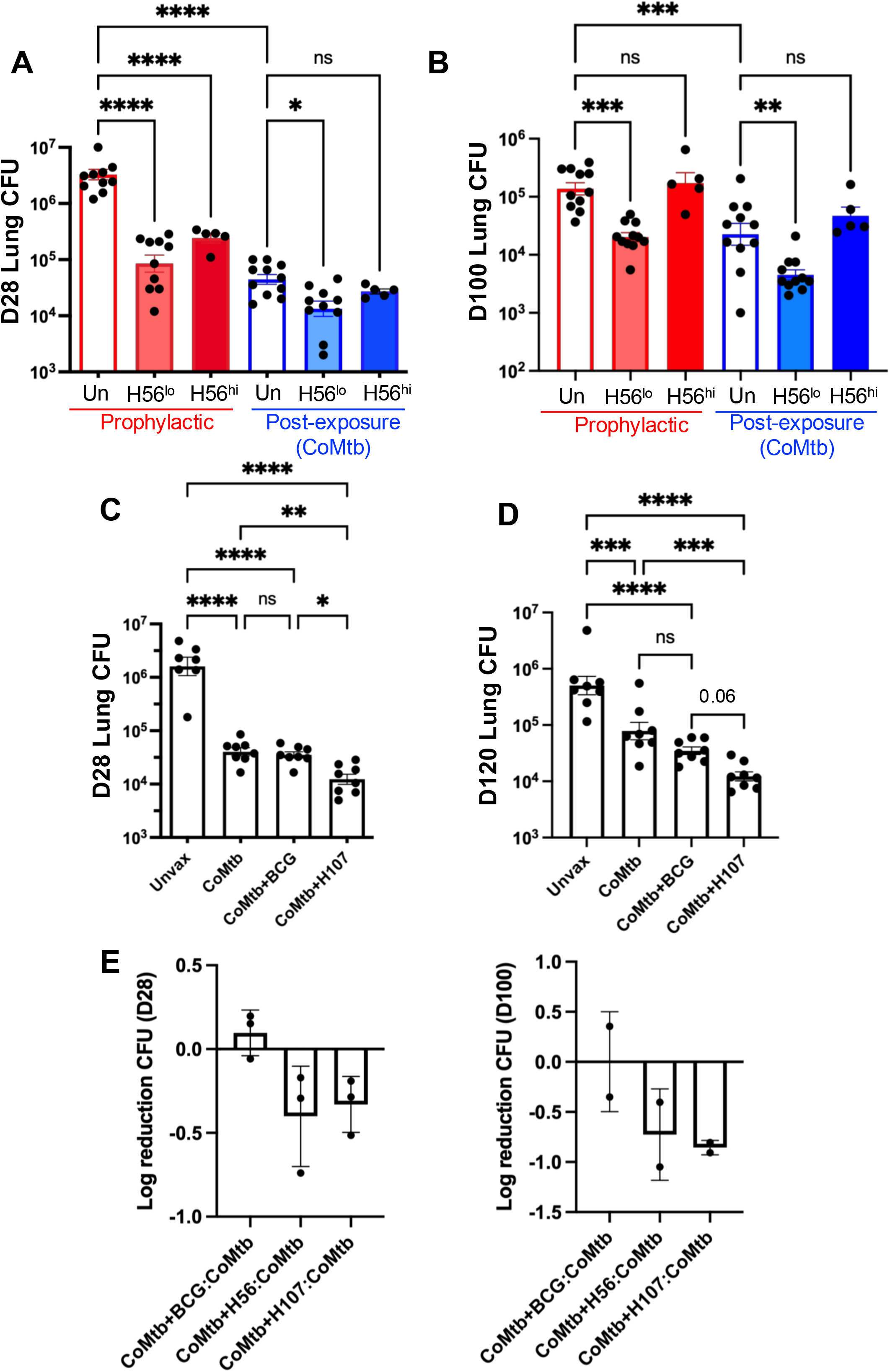
Low dose subunit vaccination is highly protective in the setting of prior Mtb exposure. (A-B) Unvaccinated and CoMtb CB6F1 mice were immunized with low (0.05 ug) or high (50 ug) doses of H56/CAF01 and subsequently aerosol-challenged with H37Rv. Lung CFU were calculated at (A) 4 weeks and (B) 14 weeks following aerosol challenge. (C-D) CB6F1 mice were left unvaccinated or administered CoMtb followed by immunization with either 10^6^ BCG or low dose (2 ug) H107/CAF01 subcutaneously. Six weeks later, mice were aerosol-challenged with H37Rv Mtb, and lung CFU were evaluated at (C) D28 post-infection and (D) D120 post-infection. (E) Cumulative data, with each dot representing an individual experiment comprising 5-8 mice per group, of log CFU reduction in lungs of mice immunized with BCG, H56/CAF01, and H107/CAF01 in Mtb-naïve or Mtb-exposed (CoMtb) settings at early (4 weeks) and late (14 weeks) times post-infection. Log-transformed CFU data are shown as the means ± SEM, and statistical analysis was performed by one-way ANOVA; *, p<0.05, **, p<0.005, ***, p<0.001, ****, p<0.0001.

We next evaluated the efficacy of H107, a different subunit vaccine currently in Phase I clinical trials (formulated with CAF01 in this study but used with CAF10b in humans(21)), using the CoMtb model and directly compared the outcomes with BCG. The H107 subunit vaccine was specifically designed to lack expression of antigens shared with BCG in order to provide synergistic protection against Mtb while preventing cross-reactivity against BCG itself(32). When CB6F1 mice were administered 0.5 μg H107/CAF01 following establishment of CoMtb, we again found that subunit vaccination provided significant additive protection over CoMtb alone at both early (Fig 2C) and late (Fig 2D) time points, whereas BCG immunization failed to improve upon the CFU reduction afforded by CoMtb. Thus, both H107/CAF01 and H56/CAF01 induced significant and sustained protection in the setting of ongoing Mtb infection (Fig 2E), whereas BCG was only protective in Mtb-naïve settings (Fig 1B).

### Decreased KLRG1+ vascular CD4 T cells is a correlate of protection of subunit vaccination in the setting of pre-existing immunity

Given the protection afforded by subunit vaccination in the setting of pre-existing immunity and the well-characterized contribution of T cells to protection(33, 34), we next sought to characterize the different phenotypes of T cells generated by vaccination in the hopes of identifying a surface marker correlate of protection. To address this, we applied unsupervised clustering to identify and quantify distinct T cell phenotypes based on surface marker expression derived from high-parameter flow cytometry profiling of lung cells collected from control and vaccinated mice 28 days post-Mtb infection using an 18-marker antibody panel (Fig 3A, Supp Fig 1A). No clusters showed increased frequency in the CoMtb+H56 group relative to all other groups. However, two clusters were found to be reduced in frequency in the CoMtb+H56 group relative to the other groups: Cluster #5 and Cluster #9 (Fig 3B, Supp Fig 1B). These two clusters were similar in their expression profiles (increased CD44, KLRG1, Ki67, Tbet, and vascular localization (IV+ staining)), except #5 represented CD8 T cells and #9 represented CD4 T cells (Supp Fig 1A). While the difference in frequencies of these unsupervised clusters did not reach statistical significance, when we validated the findings from the clustering algorithm by manually gating and quantifying the matching CD4 T cell populations, we observed significant differences (Fig 3C-F). Consistent with previous findings that H56/CAF01 immunization reduces the levels of KLRG1+ CD4+ T cell populations residing within the lung vasculature by promoting lung parenchyma homing(35), there were significantly fewer IV+ ESAT6-specific KLRG1+ CD4 T cells in H56/CAF01-immunized mice relative to unvaccinated and CoMtb mice (Fig 3C,E). This population was further reduced by H56/CAF01 or H107/CAF01 immunization in the setting of pre-existing Mtb immunity (Fig 3C,E), although the differences were not statistically significant compared to H56/CAF01 immunization in Mtb-naïve mice. In contrast, IV+ CD44^hi^KLRG1+ CD4 T cells were significantly reduced in CoMtb+H56 mice compared with all other groups, including H56/CAF01-immunized Mtb-naïve mice (Fig 3D,F), and CoMtb+H107/CAF01 mice showed a similar trend. Despite emerging as a possible distinguishing cluster by Seurat analysis, IV+ KLRG1+ CD8 T cells were not significantly different between Mtb-naïve and post-exposure H56/CAF01-vaccinated mice upon manual FlowJo gating (Supp Fig 1C), and importantly, BCG did not drive similar changes in vascular KLRG1+ CD4 T cells for either ESAT6-specific (Fig 3E) or CD44^hi^ populations (Fig 3D,F). Together, these data suggest that loss of vascular KLRG1+ CD4 T cells may be a correlate of protective responses, at least for these subunit vaccines, and indeed, KLRG1 expression levels among IV+ activated T cells significantly correlate with lung bacterial burdens (Supp Fig 1D).

**Figure 3.**
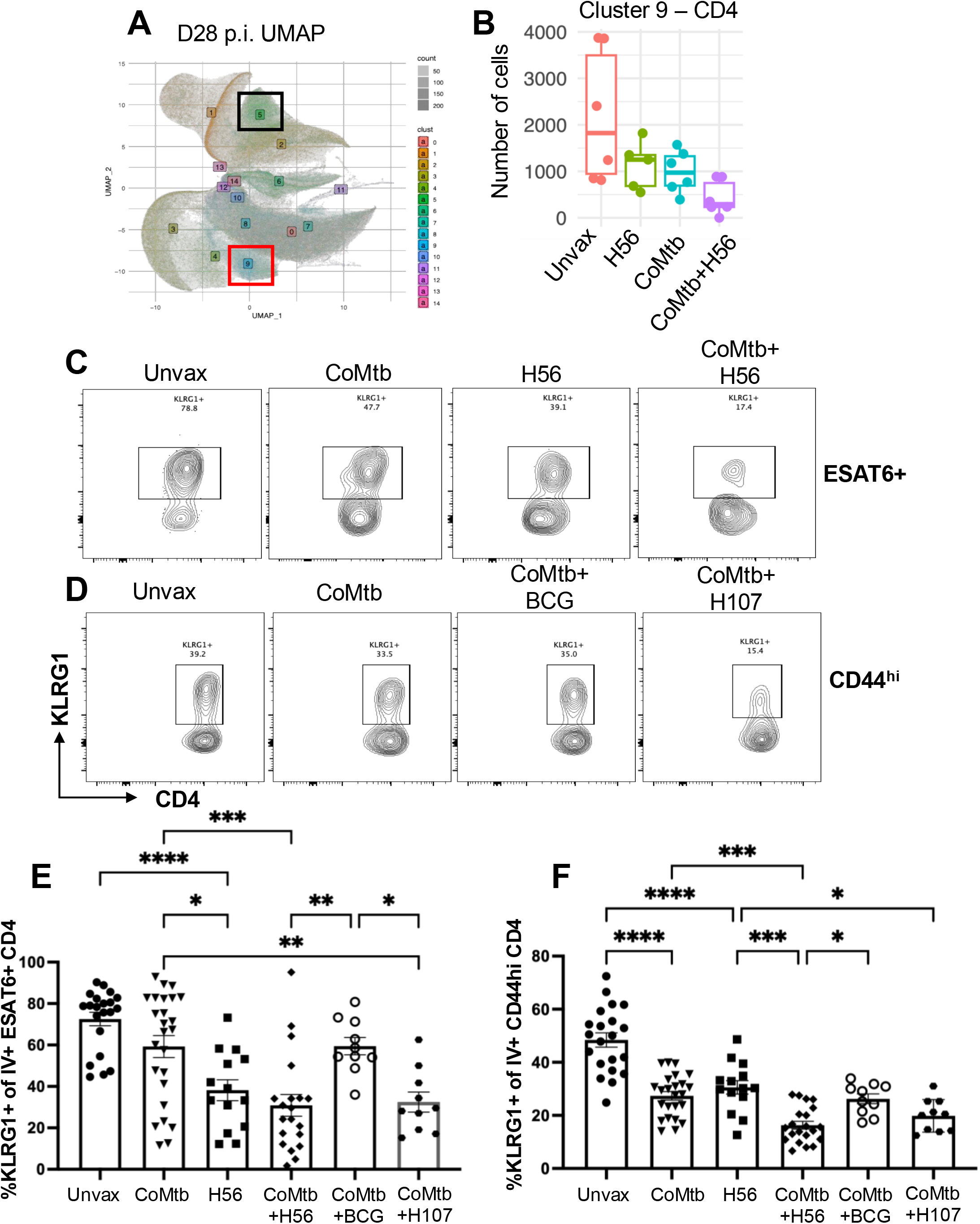
Unsupervised clustering reveals loss of KLRG1 expression on vascular CD4 T cells as a correlate of protection. (A) UMAP plot showing results of unsupervised clustering of cells isolated from Day 28 post-Mtb infection lungs applied to continuous flow-cytometry fluorescence values for 15 protein markers, summarized in 200 (X) x 200 (Y) hexagonal regions. Color intensity of each hexagonal bin of the plot is proportional to the number of cells in that region, with more opaque regions having more cells. Color shade of each region indicates the cell clusters contained in the hexagonal bin. Labels for each cluster were placed at the central (mean) point for all cluster cells. Cluster 5 (black box) represents CD8 T cells, and cluster 9 (red box) represents CD4 T cells. (B) Box-and dotplot showing total counts for cells in cluster 9, stratified by vaccination status. (C-D) Changes in KLRG1 expression on (C) vascular ESAT6-specific CD4 T cells and (D) vascular CD44 high CD4 T cells were independently validated on lung cells from mice in the different conditions 28 days following aerosol challenge. (E-F) Cumulative data on KLRG1 expression across groups among (E) vascular ESAT6-specific CD4 T cells and (F) vascular CD44 high CD4 T cells. Data are shown as the means ± SEM. Statistical analysis was performed by one-way ANOVA; *, p<0.05, **, p<0.005, ***, p<0.001, ****, p<0.0001.

### Subunit vaccination, but not BCG, drives polyfunctional CD4 T cell responses in Mtb-exposed mice

We next sought to characterize the functionality of parenchymal lung CD4 T cells generated by subunit vaccination in the setting of pre-existing immunity. To accomplish this, single cell suspensions collected from the lungs of mice administered CoMtb either alone or in combination with BCG or H107/CAF01 immunization were stimulated ex vivo with ESAT6 peptide. Then, IFNγ, IL-2, TNF, and IL-17 responses were measured by intracellular cytokine staining. Using a Boolean “OR” gating strategy, we found that significantly more CD4 T cells in the CoMtb+H107 group produced at least one cytokine compared to CoMtb alone or CoMtb+BCG (Fig 4A). This effect was likely subunit vaccine-driven, as cells stimulated with TB10.4 peptide, which is expressed by both Mtb and BCG but is absent in the vaccine formulation, showed no significant changes in cytokine-producing capacity across groups (Fig 4A). To obtain a broader look at the overall variation in cytokine production across the groups, we performed a principal component analysis (PCA) of cytokine-positive CD4 T cells and found that the CoMtb+H107 group clustered distinctly from the other 3 groups, suggesting that cells in this condition expressed a unique cytokine signature, with PC1 driven by IL-2 production (Fig 4B). We next used Boolean “combination” gating to identify all populations of cytokine-producing cells (based on production of IFNγ, TNF, and IL-2) that were induced by vaccination and therefore contributed to the unique clustering. Particularly striking was that more than half of the cytokine-producing cells in the CoMtb+H107 group made IL-2 (Fig 4C,D), consistent with a known role of these subunit vaccines in generating self-renewing, memory-like CD4 T cell populations(7, 18, 19). In contrast, only approximately one quarter of cytokine-producing lung CD4 T cells (Fig 4C,D) and <1% of total IV-CD4 T cells (Fig 4E) in the CoMtb+BCG group expressed IL-2. When the CoMtb groups immunized with either H107/CAF01 or BCG were compared, the H107/CAF01 group had significantly higher levels of triple IFNγ+TNF+IL-2+-producers and double TNF+IL-2+-producers (Supp Fig 2A) compared to the BCG group. IL-17 was only expressed at detectable levels in the CoMtb+H107 group, with statistically significant increases in parenchymal CD4 T cells that expressed IL-17 in combination with TNF or IL-2/TNF compared to the other groups (Fig 4F). These data highlight that H107/CAF01 subunit immunization confers a significant degree of polyfunctional cytokine-producing capacity in the setting of prior Mtb infection that is not recapitulated by BCG immunization.

**Figure 4.**
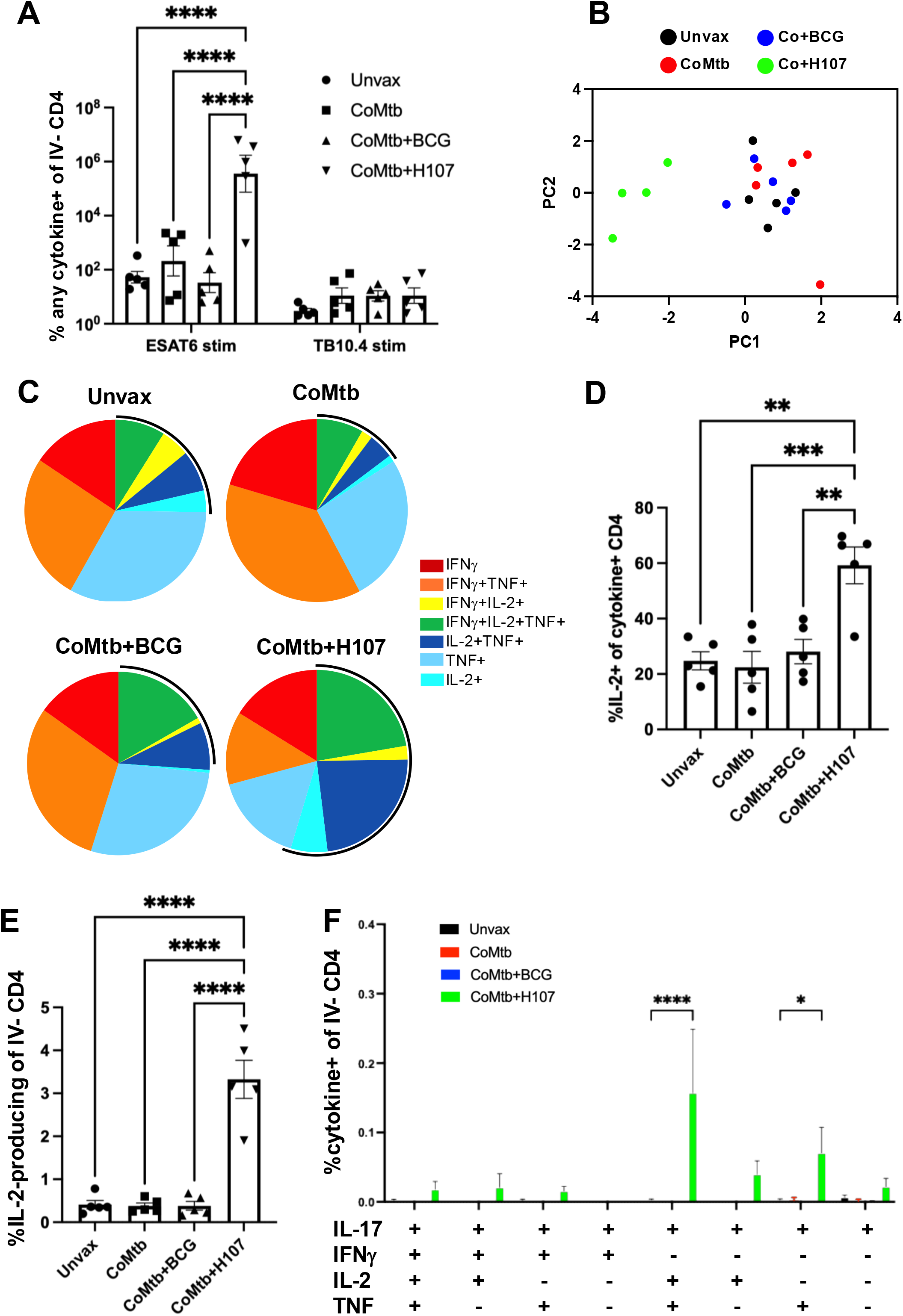
Improved protection mediated by subunit vaccination in the setting of prior Mtb exposure is associated with polyfunctional cytokine responses. CB6F1 mice were either left unvaccinated or immunized with BCG or H107/CAF01 6 weeks following intradermal Mtb infection (CoMtb). All mice were aerosol challenged with H37Rv, and lungs were harvested for flow cytometric analysis 28 days post-infection. (A) Lung cells of mice in each group were stimulated with ESAT6 or TB10.4 (p8) peptide ex vivo, and cytokine-producing capacity (positive for IFNγ OR IL-2 OR TNF OR IL-17) was evaluated among parenchymal (IV-) CD4 T cells. (B) Principal component analysis of parenchymal (IV-) lung CD4 T cells based on their capacity to produce unique cytokine combinations upon ESAT6 peptide stimulation. (C) Polyfunctional cytokine production of parenchymal CD4 T cells responding to ESAT6 peptide from each group is displayed as a pie chart, with the black arcs representing groups of cells that produce IL-2. (D) Frequency of IL-2-producing T cells among all cytokine-positive parenchymal CD4 T cells. (E) Frequency of IL-2 producing T cells among all parenchymal CD4 T cells. (F) Frequencies of distinct combinations of ESAT6-specific cytokine-producing parenchymal CD4 T cells, including IL-17, are compared across groups. Data are shown as the means ± SEM. Statistical analysis was performed by one-way ANOVA; **, p<0.005, ***, p<0.001, ****, p<0.0001.

### Subunit vaccination in the setting of CoMtb endows CD4 T cells with both memory and effector features

We next considered whether this imprinting of polyfunctionality on lung CD4 T cells was completely vaccine-driven or whether prior Mtb exposure impacted this response. To test this, we repeated the experiment with the addition of a vaccination group in Mtb-naïve mice (H107/CAF01 alone) and measured lung CFU and cytokine profiles at ∼100 days post-aerosol challenge to additionally address whether alterations in T cell function and protection were long-lived. As observed previously, prior Mtb exposure reduced lung bacterial burdens following aerosol challenge even at late timepoints, as did H107/CAF01 subunit vaccination (Fig 5A). Strikingly, H107/CAF01 immunization in the setting of prior Mtb exposure also significantly reduced lung bacterial burdens compared to immunization in the Mtb-naïve setting, whereas BCG provided no further protection over CoMtb alone (Fig 5A). Using the same gating strategy as in Fig 4A, we found that the overall cytokine-producing potential by parenchymal (IV-negative) CD4 T cells was equivalent between H107/CAF01-immunized mice in the Mtb-exposed or Mtb-naïve settings, wherein H107/CAF01 drove a dramatic increase in cytokine production in both settings (Fig 5B). Similarly, the proportion of cytokine-producing CD4 T cells that made IL-2 in response to ESAT6 stimulation was equivalently expanded in both H107-immunized groups of mice (Fig 5C). Therefore, H107/CAF01 is highly effective at priming and/or expanding memory-like CD4 T cells in both Mtb-naïve and post-exposure settings. We next examined overall differences in populations of cytokine-producing parenchymal CD4 T cells and confirmed that CoMtb predominantly maintains the effector-like features observed in the unvaccinated setting, with high frequencies of IFNγ+TNF+-producing CD4 T cells, while subunit vaccination skews the cytokine landscape towards memory-like IL-2-producing CD4 T cells (Fig 5D). Despite equivalent overall IL-2-producing CD4 T cells in Mtb-exposed or Mtb-naïve settings (Fig 5C), we found that prior Mtb exposure endowed a significant increase in vaccine-induced IFNγ expression among the IL-2-producing CD4 T cells (Fig 5E), suggesting that the cells expressed a combination of memory and effector characteristics; this finding was reproduced in an independent experiment using H56/CAF01 vaccination (Supp Fig 2B). Furthermore, the increased levels of IFNγ+IL-2+TNF+ triple-producers in response to ESAT6 stimulation, but not TB10.4 stimulation, approached statistical significance in the CoMtb+H107 group (Fig 5F). At this later timepoint, IL-17 expression was uniquely expressed by T cells in H107/CAF01-immunized mice, and the frequencies of cells were largely unaffected by CoMtb, indicating the dependence on subunit vaccination for IL-17 expression (Supp Fig 2C). Overall, these findings demonstrate that increased protection against Mtb lung burden is correlated with the generation of parenchymal CD4 T cells that share functional features of both proliferative memory T cells and terminal effectors.

**Figure 5.**
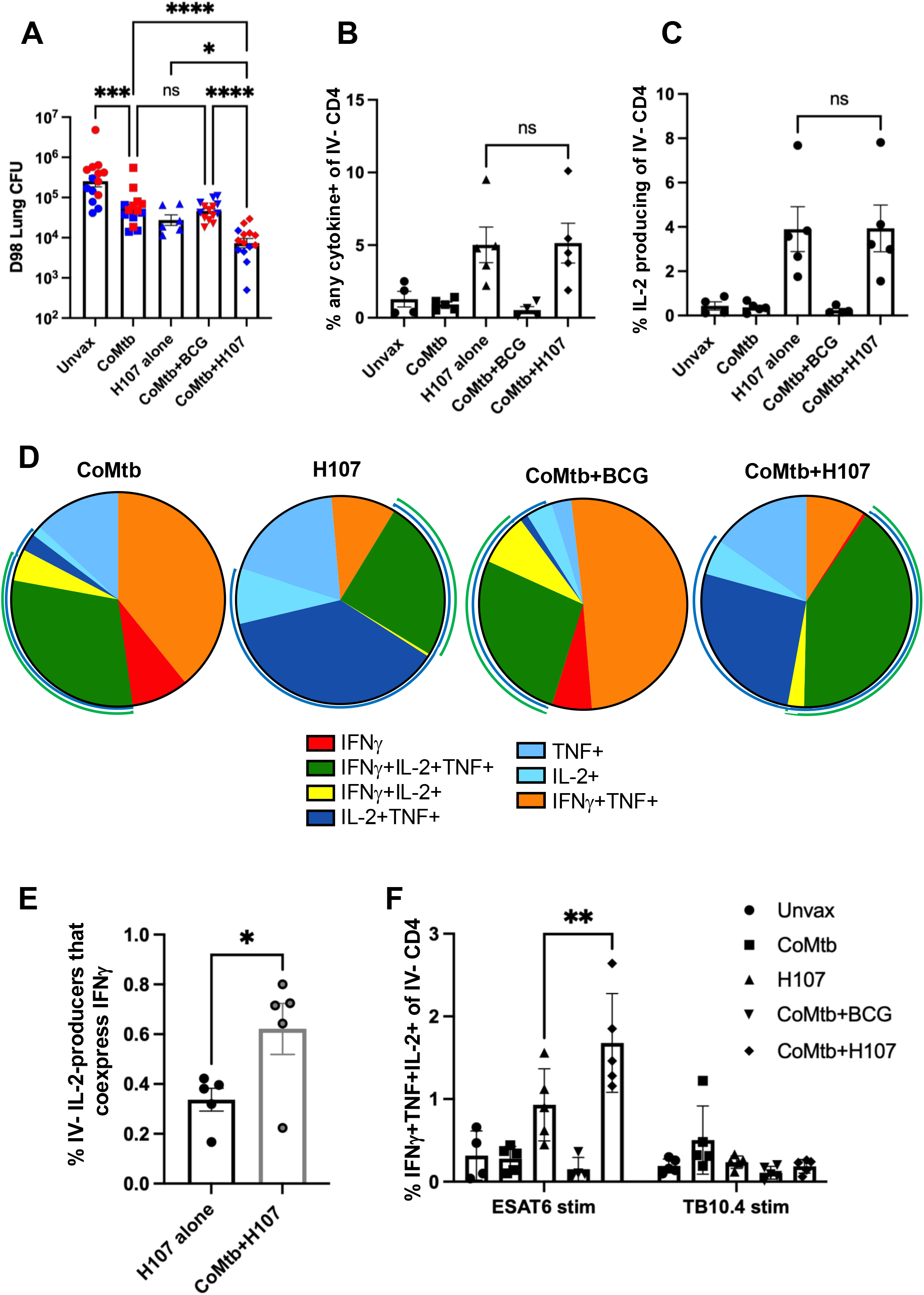
Polyfunctional cytokine responses driven by subunit vaccination are durable in the post-exposure setting. CB6F1 mice were either left unvaccinated, subunit-vaccinated, or administered intradermal Mtb (CoMtb) followed by BCG or subunit vaccination and then aerosol challenged with H37Rv for tissue collection at ∼100 days post-infection. (A) Log-transformed lung CFU at ∼100 days post-infection. Each color represents an independent experiment. (B) Cytokine-producing capacity (positive for IFNγ OR IL-2 OR TNF OR IL-17) and (C) IL-2-producing capacity were evaluated among parenchymal (IV-) CD4 T cells. (D) Polyfunctional cytokine production of parenchymal CD4 T cells responding to ESAT6 peptide from each group is displayed as a pie chart, with the blue arc representing cells that produce IL-2 and the green arc representing cells that coproduce IL-2 and IFNγ. (E) Frequency of IL-2-producing parenchymal CD4 T cells that also express IFNγ in H107-immunized mice in Mtb-naïve compared to Mtb-exposed settings. (F) Frequencies of IFNγ+TNF+IL-2+ triple-producing parenchymal CD4 T cells in response to either ESAT6 or TB10.4 peptide stimulation. Data are shown as the means ± SEM. Statistical analysis was performed by one-way ANOVA (A-C), Student’s t-test (E) or two-way ANOVA (F); *, p<0.05, **, p<0.005, ***, p<0.001, ****, p<0.0001.

### Post-exposure subunit vaccination augments Mtb-specific responses even prior to aerosol challenge

We next asked whether subunit vaccination in the setting of CoMtb leads to reprogramming of the T cell response even prior to aerosol challenge with Mtb. Mice underwent the same vaccine regimen as in prior experiments, and spleens were harvested 6 weeks following the last vaccine dose for ex vivo ESAT6 peptide stimulation. Consistent with post-challenge responses, we found that mice immunized with BCG, CoMtb or a combination of both displayed effector-like responses dominated by IFNγ, while H56/CAF01-immunized mice skewed towards IL-2 and TNF responses (Fig 6A). In striking contrast, however, mice immunized with H56/CAF01 in the setting of CoMtb showed a dramatic increase in overall cytokine abundance, with representation of both IL-2/TNF-producing and IFNγ-producing CD4 T cell subpopulations (Fig 6A). Meanwhile, CD4 T cell responses to TB10.4 peptide were largely similar across groups, with CoMtb groups expressing a small increase in IFNγ/TNF signal over Mtb-naïve mice but with no groups substantially rising above background levels (Fig 6B), suggesting that the enhanced responses to ESAT6 peptide are vaccine-driven and not reproduced by BCG. Similar to observations made following aerosol challenge (Fig 5E), when we quantified the proportion of IL-2-producing CD4 T cells that also made IFNγ, we found that mice that had been H56/CAF01-immunized following Mtb exposure had significantly more IL-2/IFNγ coexpressing CD4 T cells (Fig 6C). Consistently, there was a dramatic increase in the proportion and total numbers of IFNγ+IL-2+TNF+ triple-producers in CoMtb+H56 mice relative to all other groups, including CoMtb+BCG mice (Fig 6D, Supp Fig 3A,C) following ESAT6 stimulation but not TB10.4 stimulation (Supp Fig 3B). Because BCG does not express ESAT6 and thus would not be expected to prime a response to ESAT6 peptide, we also stimulated spleen cells with Ag85B(P25) peptide, which is present in both BCG and the H56 vaccine. As was observed with TB10.4 peptide stimulation (Fig 6B), BCG immunization in the setting of prior Mtb exposure again failed to increase either the magnitude or quality of the CD4 T cell response to Ag85B(P25) peptide relative to CoMtb or BCG alone (Supp Fig 3D), whereas the CoMtb+H56 group had increased IL-2/IFNγ co-expressing CD4 T cells (Supp Fig 3E), similar to that observed following ESAT6 stimulation both pre-challenge (Fig 6A,C) and post-challenge (Fig 5E). Interestingly, the magnitude of the response to Ag85B(P25) stimulation was similar between the Mtb-naïve and post-exposure H56/CAF01 groups, suggesting a difference in the response profiles to ESAT6 and Ag85B, as has been reported previously(36) (Supp Fig 3D).

**Figure 6.**
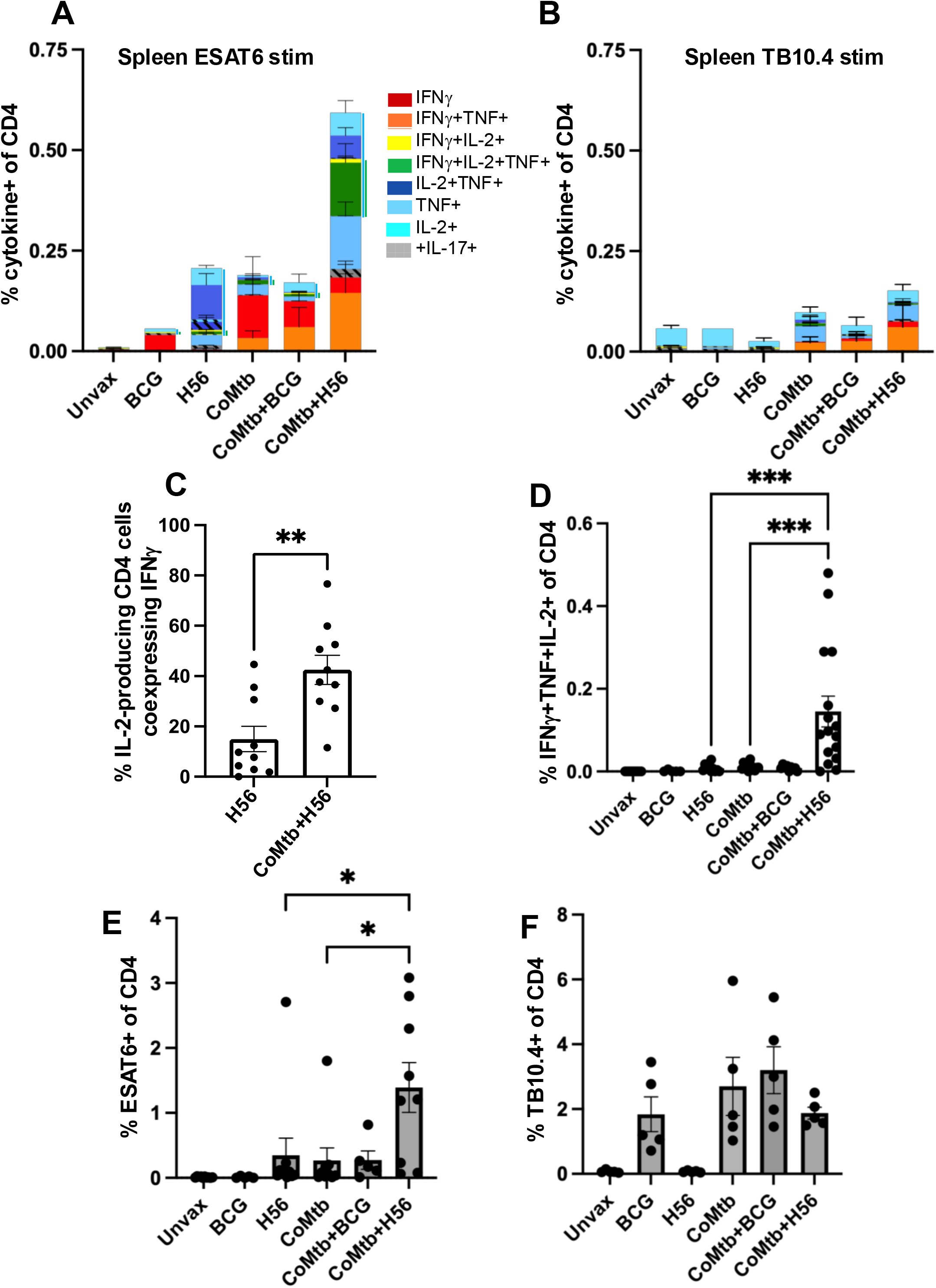
Vaccine-specific immune responses are augmented by Mtb exposure prior to aerosol challenge. (A-F) CB6F1 mice were left unvaccinated, immunized with BCG or H56, or administered intradermal Mtb (CoMtb model) followed by BCG or H56 immunization. Spleens and lungs were harvested 6 weeks following the final immunization for ex vivo peptide stimulation and tetramer analysis. (A-B) Splenocytes from mice in each group were stimulated ex vivo with (A) ESAT6 and (B) TB10.4 p8 for intracellular cytokine analysis. Shown are frequencies of overall cytokine-producing cells on the x-axis, with the relative distributions of cytokine-producing combinations shown within the bars. The blue lines adjacent to the bars in the ESAT6 graph represent cytokine combinations that produce IL-2, and the green lines indicate cell that coproduce IL-2 and IFNγ. (C) Comparison of the IL-2/IFNγ-producing capacity of splenocytes responding to ESAT6 peptide from H56-immunized mice in Mtb-naïve and Mtb-exposed settings. (D) The frequency of triple-producing IFNγ+TNF+IL-2+ splenic CD4 T cells following ESAT6 peptide stimulation in each group of mice. (E-F) Tetramer staining for (E) Frequency of ESAT6-specific and (F) TB10.4-specific CD4 T cells in the lungs of mice from each group. Data are shown as the means ± SEM. Statistical analysis was performed by Student’s t-test (C) or one-way ANOVA (D,E); *, p<0.05, **, p<0.005, ***, p<0.001.

We further examined the levels of Mtb-specific CD4 T cells in the lungs to determine if the immune landscape at the future site of infection was altered by vaccination prior to challenge. We observed differential expansion of Mtb-specific CD4 T cells that, importantly, were unlikely to be induced directly by colonization of the lung by CoMtb-derived bacteria because we were never able to culture Mtb from these lungs. ESAT6-specific CD4 T cells could be detected in each group that had been immunized with an ESAT6-containing regimen, either H56/CAF01 or Mtb itself, but unvaccinated mice or those receiving only BCG did not have detectable ESAT6-specific T cell populations, as expected since BCG does not express ESAT6 (Fig 6E, Supp Fig 4E). Interestingly, mice in the CoMtb+H56 group had significantly increased levels of ESAT6-specific lung CD4 T cells relative to either H56/CAF01-immunized or CoMtb alone mice, indicating that lung cells, in addition to peripheral splenic cells, were also expanded by vaccination prior to aerosol challenge (Fig 6E, Supp Fig 4E). Consistent with their functional phenotype in the spleen, these ESAT6-specific CD4 T cells in the lung parenchyma of CoMtb+H56 mice also produced significantly higher levels of IFNγ/IL-2/TNF compared to the vaccine or CoMtb alone groups (Suppl Fig 4A,B,D). In contrast, TB10.4-specific lung CD4 T cells were equally expanded both in terms of cytokine-production (Supp Fig 4C) and tetramer-binding (Fig 6F, Supp Fig 4F) in all groups of mice receiving BCG or Mtb-containing vaccine modalities, consistent with the splenocyte data that the enhanced T cell expansion is subunit vaccine-dependent (Fig 6B,F). Together, these data show that subunit vaccination in the setting of prior Mtb exposure strongly expands populations of polyfunctional cytokine-producing CD4 T cells, including lung tissue-resident cells, even prior to aerosol exposure, which may contribute to the enhanced protection observed in the lungs of CoMtb+H56 and CoMtb+H107 mice following aerosol challenge.

### Cytokine-expressing T cells from QFT+ H56-immunized humans recapitulate findings in CoMtb-immunized mice

We next asked whether our findings in CoMtb mice could be observed in vaccinated humans with prior Mtb exposure. To address this, we analyzed data from a cohort of QFT-(n=15) and QFT+ (n=12) adults who were either unvaccinated or immunized with two 5 µg doses of H56/IC31 as part of the H56-035 clinical trial performed at the South African Vaccine Initiative (SATVI) in collaboration with the Statens Serum Institut (SSI). Blood was collected at enrollment (baseline) as well as at a memory time point following vaccination (224-292 days post-dose 1), and PBMC were stimulated ex vivo with an ESAT6 peptide pool, complementing our mouse assays of splenocytes harvested from Mtb-naïve (i.e., QFT-) or CoMtb (i.e., QFT+) mice with or without H56/CAF01 immunization, as described in Figure 6. We found that frequencies of cytokine-expressing CD4 T cells in blood from individuals with Mtb infection alone (QFT+ baseline) were higher than those observed following vaccination (QFT-memory). However, vaccination in the setting of prior Mtb exposure (QFT+ memory) led to a further significant increase in frequencies of cytokine-expressing CD4 T cells relative to either Mtb-naïve/immunized or Mtb-exposed/unimmunized individuals (Fig 7A). This was similar to the findings in mice shown in Fig 6A, and replotted to match the human cohorts in Fig 7B, wherein splenic CD4 T cells from the CoMtb+H56 group demonstrated the highest cytokine-producing potential of all groups, with higher levels than CoMtb or H56/CAF01 alone. We further observed strong concordance between the human and murine data with both showing a significant enhancement of IL-2 producing CD4 T cells that co-express IFNγ (Fig 7C; see also Fig 6C) as well as IFNγ+TNF+IL-2+ triple-producing CD4 T cells (Fig 7D; see also Fig 6D) elicited by vaccination in the setting of prior Mtb exposure. Finally, we observed that, among QFT+ individuals, the immune response elicited by higher vaccine dose was reduced relative to the lower dose (Fig 7E), mirroring our findings in mice (Fig 2B). Overall, the presence of prior Mtb infection, either based on QFT positivity in humans or CoMtb infection in mice, confers an enhanced polyfunctional cytokine response to subunit vaccination that is correlated with protection against challenge in the murine TB infection model. Our findings suggest that subunit vaccines can induce robust immune responses in both mice and humans despite prior Mtb exposure. In mice these responses were associated with enhanced protection, supporting the potential of subunit vaccines over live attenuated vaccines, to address the longstanding goal of curbing TB disease and transmission among adolescents and adults in endemic settings.

**Figure 7.**
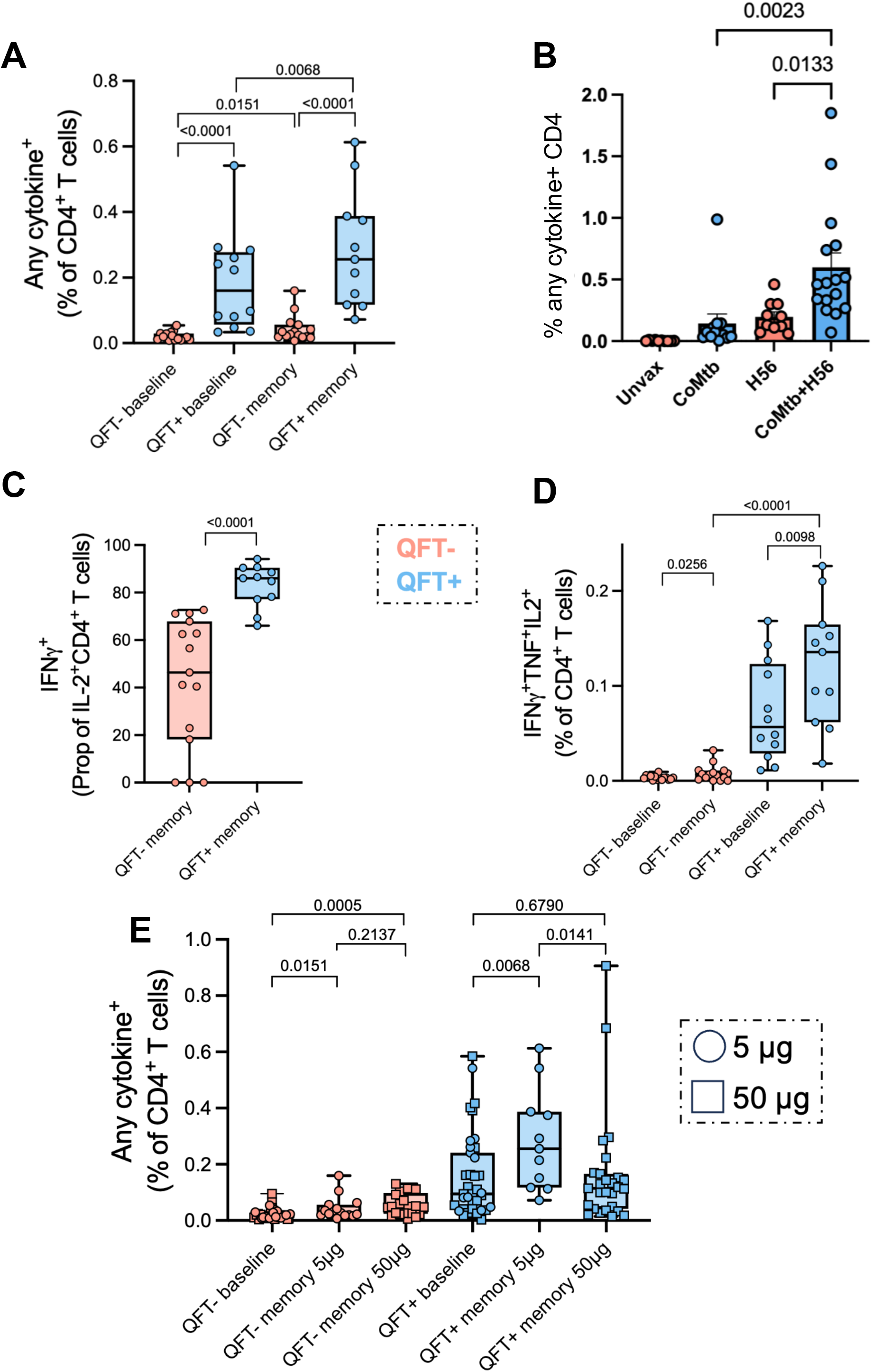
Human clinical trial PBMC cytokine data recapitulate findings in the CoMtb mouse model. As part of the H56-035 clinical trial performed at SATVI in collaboration with SSI(26), 15 QFT-adults and 12 QFT+ adults were immunized with two doses of 5 ug H56/IC31 2 months apart, and whole blood was collected at baseline and again at a memory time point (∼8 months after the last vaccination) for ESAT6 stimulation to measure cytokine responses. (A) Frequencies of peripheral blood ESAT6-specific CD4 T cells capable of producing any combination of IFNγ, IL-2, TNF and/or IL-17 (any cytokine) was measured in QFT-and QFT+ participants before (baseline) and after (memory) H56/IC31 immunization. (B) The same murine splenocyte data from Fig 6A is replotted to directly compare to the human PBMC data. (C) Proportions of ESAT6-specific peripheral blood IL-2+ CD4 T cells that co-express IFNγ in QFT-and QFT+ humans immunized with H56/IC31. (D) Frequencies of ESAT6-specific triple-producing IFNγ+TNF+IL-2+ peripheral blood CD4 T cells in QFT-and QFT+ individuals before and after H56/IC31 vaccination. (E) Frequencies of ESAT6-specific CD4 T cells in participants receiving two doses of 5 or 50 ug of H56/IC31. Data in (E) combine results from the H56-035 trial described above (5ug dose) and the H56-032 trial(23), which included 27 QFT+ and 21 QFT-participants who received 50 ug of H56/IC31 2 months apart and whole blood was collected at baseline and at a memory time point (6 months after the last vaccination). Cross-sectional study groups were compared by Mann-Whitney test and paired longitudinal data were analysed by Wilcoxon test. Unadjusted p values are shown.

## Discussion

This study sought to establish a murine model for vaccine efficacy testing that incorporates pre-existing immunity to mycobacteria, an important consideration given that clinical trials in humans currently focus on adolescent and adult populations living in TB-endemic regions who may be frequently exposed to Mtb. Here we used a chronic Mtb lymph node infection (CoMtb) to establish pre-existing Mtb immunity, a model that has been shown to induce concomitant immunity to an autologous aerosol Mtb challenge (10, 37). In this setting of prior Mtb immunity, we found that the adjuvanted subunit vaccines H56/CAF01 and H107/CAF01, but not BCG, were able to significantly improve upon the protection afforded by CoMtb alone.

We attempted to identify correlates of protection related to H56/CAF01 immunization in the setting of pre-existing immunity to Mtb. We first noted that, even prior to Mtb aerosol challenge, significantly higher frequencies of ESAT6-specific cytokine-producing CD4 T cells were induced in the spleens of subunit-vaccinated CoMtb mice relative to the mice receiving either vaccine or CoMtb alone, suggesting the presence of a heightened memory response. In contrast, BCG immunized mice did not display any boosting in the setting of CoMtb, either in the magnitude or quality of cytokine-expressing T cells, following stimulation with Ag85B antigen, which is present in both BCG and H56. H56-immunized mice, however, showed an enhanced polyfunctional response of IL-2/IFNγ co-producers, as observed in the lung following Mtb challenge. In general, we found that subunit vaccination was associated with the induction of polyfunctional immune responses highlighted by the presence of IL-2 and IL-17, consistent with many prior reports using CAF-containing vaccines and with the notion that polyfunctional and IL-17 producing T cell responses are strong predictors of Mtb control in mice(5, 18, 20, 38, 39). This is likely also true in humans, as increasing bacterial loads in humans inversely correlated with IFNγ+TNF+IL-2+ CD4 T cells, a pattern that was reversed upon antimicrobial treatment(40), the correlation of IL-17 responses with resistance to TB progression(41, 42), and the detection of Th17 cells in BCG-immunized NHP(43, 44). Studies both in NHP and humans have previously shown that these polyfunctional cytokine responses are enhanced by low-dose subunit vaccination in settings of previous Mtb infection, consistent with our data in the CoMtb murine model. For example, higher vaccine-induced T cell responses were observed among QFT+/Mtb-exposed individuals in the setting of ID93/GLA-SE(45), H1:IC31(24), M72/AS01E(46, 47), and even H56:IC31(23) vaccinations. The concordance of these observations between human and mice support the use of the CoMtb model to measure vaccine-mediated cytokine responses associated with protection in previously Mtb-infected humans and provide a novel opportunity to correlate these responses with protection in the mouse.

A reliable correlate of protection against TB has yet to be identified, and the induction of polyfunctional T cells is not always indicative of vaccine success(48). This may be in part because the capacity to produce cytokines does not guarantee that T cells are properly localized to the site of infection. Hence, other factors, such as self-renewal capacity(49, 50), T cell homing(19, 35, 38), antigen recognition(36, 51, 52), and exhaustion(53–55), can also contribute to protective T cell features. There is evidence in non-human primates, however, that polyfunctional T cells localized properly do correlate with protection against TB in animals vaccinated intravenously with a high dose of BCG, suggesting that lung localized, polyfunctional T cells may still represent a more general protective feature that can be extended beyond mouse models(56, 57). In future studies, the CoMtb model may provide a platform to dissect mechanisms of protection in the context of prior immunity, and insights from such studies could generate novel biomarkers to predict vaccine efficacy in human clinical trials.

In our study, the magnitude of the lung cytokine-expressing CD4 T cell response was largely driven by subunit vaccines irrespective of prior Mtb immunity, but the qualitative nature of the responses, as defined by the composition of immune cells co-expressing IFNγ and IL-2, was significantly augmented by vaccination in the setting of CoMtb. Combined, these data indicate that the immune landscape is altered by subunit vaccination in the setting of pre-existing Mtb immunity, providing both quantitative and qualitative improvements in cytokine-expressing T cell responses relative to unvaccinated mice. We also noted that subunit vaccination in CoMtb mice led to a significant reduction in KLRG1 expression among activated CD4 T cells in the lung vasculature. H56/CAF01 has previously been associated with loss of KLRG1+ IV+ T cells in mice, which was linked to better T cell homing to the lung parenchyma and less trapping in the lung vasculature(35). KLRG1 expression is associated with terminal differentiation and high levels of IFNγ production in both mice and humans, and when adoptively transferred into mice, this subset fails to provide protection against Mtb challenge (38, 55, 58, 59), suggesting that their propensity for being retained in the vasculature and failing to reach the lung parenchyma restricts their protective effector functions. It is tempting to speculate that perhaps subunit vaccination in the setting of prior immunity enhances lung homing of Mtb-specific CD4 effector T cells, enabling vaccine-induced T cells to better access the site of infection and reduce bacterial burdens, although future experiments will be needed to directly test this question. It remains unclear if peripheral blood T cell expression of KLRG1 could function as a biomarker of vaccine efficacy in humans, as there is discrepancy in the field as to whether KLRG1 expression is increased(58) or reduced(59) on blood T cells of active TB patients relative to healthy controls, and KLRG1 expression does not appear to restrict CD4 T cell extravasation across the lung endothelium in rhesus macaques during Mtb infection(60). Our finding of reduced KLRG1 expression in vascular CD4 T cells in the setting of protective immunity is specific to either antigen-specific or CD44^hi^ T cells, and this degree of discrimination may require further exploration within human populations.

Our findings are consistent with the long-standing observation that BCG is most effective when administered to mycobacteria-naïve infants at birth but becomes less effective when administered to school-aged children, who are more likely to have pre-existing mycobacteria-specific immunity(1, 2). There are two prevailing hypotheses, known as the masking and blocking hypotheses, as to why BCG vaccination in adolescence, and more recently BCG revaccination(61), may be ineffective at preventing TB disease (29). The former surmises that immunity resulting from the initial mycobacterial exposure masks any vaccine-induced effects that arise because the pre-existing response is already protective, whereas the latter predicts that prior immune responses prevent a new mycobacterial-based vaccine from achieving sufficient vaccine take. Our data are consistent with the blocking hypothesis, as significantly lower levels of BCG CFU were detected in CoMtb mice with prior Mtb immunity, suggesting that the prior mycobacterial responses reduced the ability of BCG to persist and replicate. This observation suggests that vaccines that are overly similar to endemic mycobacterial strains may be ineffective in already highly exposed populations. Immunologically, our data may also indicate some level of masking, as BCG was still detectable, though at lower levels, in our CoMtb-immunized mice; thus, the immune response elicited by BCG may have also been unsuccessful because it was overly similar to the CoMtb-induced response that was dominated by IFNγ and TNF without the addition of IL-2 and/or IL-17. Overall, our data suggest that the use of adjuvanted subunit vaccines in adolescents and adults may be advantageous through the capacity to de novo prime and/or boost pre-existing natural immunity, whereas immunizing already mycobacteria-exposed individuals with a live mycobacterial vaccine such as BCG may fail to provide any additional protective immunity.

A recent example of this type of success comes from the M72/AS01E phase 2b trial that showed 49.7% efficacy in prevention of disease (POD) amongst QFT+ individuals, a milestone for the TB vaccine field and underscoring the potential of subunit vaccines to provide protection in highly endemic regions(62). Immunogenicity studies for M72/AS01E showed similar results as those described in our murine study, with higher overall levels of cytokine-producing CD4 T cells in TST+ individuals than TST-individuals(47). Unfortunately, the recent prevention of recurrence (POR) trial of H56/IC31, which enrolled active TB patients treated with antibiotics prior to vaccination, failed to show efficacy. While the Phase I safety and immunogenicity data of this vaccine were encouraging(27), H56:IC31 immunization resulted in increased rates of disease relapse as compared to the placebo group; although, this difference did not reach statistical significance, further development of H56/IC31 has been suspended(22). It is important to note that our murine study was more reflective of a POD trial of QFT+ individuals, as the mice had not been previously aerosol challenged and subsequently antibiotic-treated as in the POR trial. Furthermore, vaccine outcomes can vary heavily based on the timing of vaccine administration, i.e., as a therapeutic vaccine given either at the time of treatment or post-treatment. Because H56/IC31 will never be tested in a true POD trial, it will remain unknown how predictive our animal model data would have been in the human setting, and it remains imperative that we find ways to preclinically assess vaccine candidates in a relevant model of Mtb-exposed individuals.

Despite the promise of CoMtb as a model for pre-existing immunity in the setting of vaccination, there remain limitations. QFT positivity in humans represents a large spectrum of individuals, ranging from those with asymptomatic TB (those who harbor Mtb bacilli but lack disease symptoms) to those who were previously exposed but perhaps were able to clear the infection. CoMtb, as a model, likely represents only a limited subset of that spectrum. Furthermore, it remains unclear where on the spectrum of QFT positivity lies the highest degree of susceptibility to developing disease and whether that level is fully recapitulated by the CoMtb model. In future studies, a vaccine that is known to have efficacy in QFT+ individuals, such as the M72/AS01E vaccine, should be tested in the setting of CoMtb to determine whether the protection can be reproduced. If it is not, that might suggest either that QFT+ individuals who have already cleared their Mtb infection might be the most likely to benefit from vaccination or that the QFT+ population that can be protected by vaccination is too heterogeneous to model with CoMtb. Such a scenario may be better modeled by antibiotic treating CoMtb mice prior to vaccination, and future studies could address this using new vaccine candidates.

## Materials and Methods

### Sex as a biological variable

Only female mice were used in the experiments in this study. The human cohort represented an equal mix of males and females.

### Mice

CB6F1 (C57BL/6 x Balb/c) mice were purchased from the Jackson Laboratories (Bar Harbor, ME). All mice were housed and maintained in specific pathogen-free conditions at Seattle Children’s Research Institute (SCRI).

### Concomitant Mtb (CoMtb) model

The CoMtb model was established as described previously(10). Briefly, CB6F1 mice were first anesthetized by i.p. injection of 400 ul of ketamine (4.5 mg/ml) and xylazine (0.5 mg/ml) diluted in PBS. Mice were placed in a lateral recumbent position, and the ear pinna was flattened with forceps and pinned onto an elevated dissection board using a 22g needle. H37Rv Mtb grown to an OD between 0.2-0.5 over a 48-hour period was diluted to 10^6^ CFU/ml in PBS, and 10 ul (10^4^ CFU) was administered into the dermis of the ear using a 26sg Hamilton syringe. Mice were then rested for 6 weeks prior to subsequent aerosol challenge or tissue harvest.

### Immunizations

For BCG immunizations, BCG was grown to OD 0.2-0.5 in 7H9 media supplemented with 10% OADC (Middlebrook), 0.2% glycerol (Sigma-Aldrich), and 0.05% Tween-80 (Sigma-Aldrich). BCG was then diluted to 5x10^6^ CFU/ml in PBS, and 10^6^ was administered via subcutaneous injection (s.c.) in a 200 ul volume. For H56 and H107 subunit vaccinations, protein antigen kindly provided by the Statens Serum Institut (SSI) was diluted in 10 mM Tris buffer to the appropriate concentration (0.05 ug, 0.5 ug, 2 ug, or 50 ug depending on the experiment), mixed with an equal volume of CAF01 cationic liposomal adjuvant, and vortexed for 30 seconds to fully emulsify the formulation. The vaccine was administered s.c. in a 200 ul volume every other week for 3 total injections. Mice were then rested for 6 weeks prior to aerosol Mtb challenge or tissue harvest for pre-infection analyses.

### Aerosol infections

Most aerosol infections were performed with a stock of wildtype H37Rv Mtb. To perform standard dose aerosol infections, mice were enclosed in a Glas-Col aerosol infection chamber, and ∼50-100 CFU were deposited directly into their lungs. In order to confirm the infectious dose, two mice in each infection were immediately sacrificed and their lung homogenates plated onto 7H10 plates for CFU enumeration.

### Bacterial CFU enumeration

To determine bacterial loads in tissues of Mtb-infected mice, the left lung of each mouse, and the cervical lymph node of mice receiving intradermal CoMtb, was homogenized in 0.05% Tween 80 in PBS. Then, 10-fold serial dilutions were made in 0.05% Tween 80 and plated onto 7H10 plates. Colonies were counted after 21 d of incubation at 37°C to determine CFUs per organ.

### Lung and spleen single cell suspensions

At the indicated times post-infection, mouse lungs were excised and lightly homogenized in HEPES buffer containing Liberase Blendzyme 3 (70 μg/ml; Roche) and DNaseI (30 μg/ml; Sigma-Aldrich) using a gentleMacs dissociator (Miltenyi Biotec). The lungs were then incubated for 30 min at 37°C and then further homogenized a second time with the gentleMacs. The homogenates were filtered through a 70 μm cell strainer, pelleted for RBC lysis with RBC lysing buffer (Thermo), and resuspended in FACS buffer (PBS containing 2.5% FBS and 0.1% NaN_3_). For splenocyte isolation, spleens from experimental mice were gently crushed between frosted glass slides, passed through a 70 μm cell strainer, and RBC-lysed prior to resuspension in FACS buffer.

### Surface antibody staining

Single cell suspensions were first washed in PBS and then incubated with 50 μl Zombie Aqua viability dye (BioLegend) for 10 min at room temperature in the dark. Viability dye was immediately quenched by the addition of 100 μl of a surface antibody cocktail diluted in 50% FACS buffer/50% 24G2 Fc block buffer using saturating levels of antibodies. Surface staining was performed for 20 min at 4°C using combinations of the following antibodies: CD3 (BioLegend, 100328, 100355; BD, 367-0031-82), CD4 (BioLegend, 100559; BD, 569180; Invitrogen, 25-0042-82), gdTCR (BD, 748989), CD8α (Invitrogen, MCD0830; BioLegend, 100723), NK1.1 (BioLegend, 108736), CD153 (BD, 748917, 740542, 740942), CD69 (BioLegend, 104536), PD1 (BioLegend, 135240), CD62L (Invitrogen, 47-0621-82), CD11b (Invitrogen, 56-0112-82), CD11c (Invitrogen, 56-0114-82), (KLRG1 (BD, 740279; BioLegend, 138410), CD40L (Invitrogen, 25-1541-82), SiglecF (BD, 746668), and CD44 (BD, 563971; BioLegend, 103040; Invitrogen, 47-0441-82). To stain for antigen-specific MHCII-restricted T cells, tetramer reagents either generated in-house(63) or provided by the National Institutes of Health Tetramer Core Facility were included in the antibody cocktail, and cells were incubated at room temperature for 1 hour while protected from light. Tetramers used in this study recognized ESAT6_4-17_ and TB10.4_73-88_. Then, the cells were washed once with FACS buffer and fixed with 4% paraformaldehyde for 30 min prior to analyzing on an LSRII flow cytometer (BD Biosciences) or an Aurora spectral cytometer (Cytek). In cases where vascular-resident cells were labeled in vivo, mice were deeply anesthetized with isoflurane, and 1 ug of APC-conjugated antibody against CD45.2 (BioLegend, 109814) or CD4 (BioLegend, 116014) was infused intravenously 5-10 min prior to sacrifice.

### Ex vivo peptide stimulations and intracellular cytokine staining

Single cell suspensions from lungs and/or spleens were prepared in complete RPMI growth media (RPMI 1640 supplemented with 10% FCS, 2 mM l-glutamine, 10 mM HEPES, 0.5 µM 2-ME, 100 U/ml penicillin, and 100 µg/ml streptomycin containing), and approximately 10^6^ cells were plated into round-bottomed 96-well plates in the presence of 5 ug/ml peptide [ESAT6_4-17_ (amino acid sequence: QQWNFAGIEAAASA), TB10.4_71-88_ p8 (amino acid sequence: AMSSTHEANTMAMMARDT), or Ag85B_240-254_ (amino acid sequence: FQDAYNGAGGHNAVF)] or 5 ug/ml protein antigen (H56 or H107), 1 ug/ml each anti-CD49d/anti-CD28, and 10 ug/ml Brefeldin-A. Cells were incubated at 37°C in a 5% CO_2_ incubator for 4-5 hours and surface-stained with the indicated antibody cocktails as described above. Then, cells were fixed with either the Invitrogen fixation/permeabilization kit (Invitrogen) or the BD Cytofix/Cytoperm kit (BD) overnight at 4°C. The following day, cells were permeabilized with the appropriate permeabilization wash buffer and incubated with intracellular antibodies against IFNγ (BioLegend, 505817), TNF (AF700, clone: MP6-XT22, BD), IL-2 (BioLegend, 503808), and IL-17 (BioLegend, 506298) diluted in permeabilization buffer for 20 min at 4°C. In other experiments, intracellular staining was performed for Ki67 (BioLegend, 652413), T-bet (Invitrogen, 45-5825-82), Rorgt (Invitrogen, 53-6981-82), and Foxp3 (Invitrogen, 48-5773-82). Cells were then washed once with FACS buffer and analyzed on an LSRII flow cytometer (BD Biosciences) or an Aurora spectral cytometer (Cytek).

### Unsupervised clustering of flow cytometry data

Unsupervised clustering of flow cytometry data was performed using the R ggycto(64) and Seurat(65) packages. Flow-cytometry data was loaded from .fcs files and initial gating was perfomed with ggcyto to retain CD3+ single-cell lymphocytes. Each sample was randomly downselected to 48,000 cells to match the least abundant sample and ensure a directly comparable total number of cells across each sample. Fifteen protein markers were used to perform unsupervised clustering: IV CD45.2, PD1, Rorgt, KLRG1, CD4, gdTCR, Ki67, CD44, CD153, ESAT6, CD40L, CD69, CD8, T-bet, and Foxp3. Bi-exponential scaling was then applied to each marker fluorescence value, and Seurat was used to scale, center, PCA transform and run unsupervised clustering on the data, using the ‘igraph’ clustering method. Cluster counts for each unsupervised cluster were then quantified for downstream analysis. UMAP was used to visualize clusters, using Seurat to calculate UMAP coordinates based on PCA-transformed data.

### Human studies

Analysis of human data combined QFT-and QFT-study participants enrolled in specific arms of the H56-032 (two doses of 50 ug H56/IC31)(23) and H56-035 (two doses of 5 ug of H56/IC31)(26) clinical trials. As described in the original publications, fresh whole blood was stimulated with peptides pools spanning the H56 antigens, including ESAT-6 and analysed by flow cytometry using similar panels of antibodies. For this study, raw data was re-gated in FlowJo using a harmonized gating strategy, exported, integrated and further analysed in R.

### Statistics

All murine data are shown as the means plus standard error of the mean. Data within individual experiments were compared by unpaired two-tailed Student’s t-test when only two groups were involved. Experiments with more than two groups were compared by one-way ANOVA with Tukey’s multiple comparison test or two-way ANOVA. p<0.05 was considered statistically significant, with * indicating p<0.05, ** indicating p<0.005, *** indicating p<0.001, and **** indicating p<0.0001. For human data, box plots showing the frequencies of CD4 T cells are shown. Cross-sectional study groups were compared by Mann-Whitney test, and paired longitudinal data were analysed by Wilcoxon test. Unadjusted p values are shown.

### Study Approval

For all murine studies performed at SCRI, approval was granted by the IACUC under approval number #499. All human studies were conducted according to the Declaration of Helsinki principles, as previously described(23, 26).

## Data Availability

Murine and human data are available in the Supporting Data Values file.

## Author Contributions

SBC, JSW, TL, RM, and KBU contributed to conceptualization. JSW, TL, RM, and KBW contributed to funding acquisition. SBC performed the experiments. SBC, FJD, JDA, EN, and AG contributed to analysis. JSW, TL, TJS, EN, RM, and KBU contributed to supervision. SBC wrote the original draft. All authors participated in reviewing and editing, and all authors approved the final manuscript.

## Funding Support

This project was supported by an NIH grant awarded to KBU and TL (R01AI134246) and an NIH IMPAc-TB contract awarded to KBU (75N93019C00070).

## Acknowledgments.

We would like to thank Peter Andersen, Courtney Plumlee, and Lauren Cross for helpful discussions. We acknowledge technical support by Suk-Lin Zhou, Bridget Alexander, and Lindsay Engels. We thank the NIH Tetramer Core facility (NIH Contract 75N93020D00005 and RRID:SCR_026557) for providing ESAT6 and TB10.4 tetramer reagents. We are thankful to the clinical trial participants, the SATVI clinical and laboratory teams and Ms Kelly Williams for statistical support.

## Conflict of Interest

The authors have declared that no conflict of interest exists.

**Supplemental Fig 1. Unsupervised clustering of lung T cells.** (A) Heatmap showing median fluorescent intensity for protein surface markers for each Seurat-determined unsupervised cluster. Median intensities were plotted after scaling and centering (to mean 0 and stdev 1) per marker. Hierarchical clustering was used to determine x-axis cluster order and y-axis marker order. (B) Box and dotplot showing total counts for cells in cluster 5 stratified by vaccination status. (C) Frequency of KLRG1 expression among CD8 T cells that match the Seurat phenotype for cluster 5 (CD44+IV+CD8+). (D) Correlation between KLRG1 expression among IV+CD44+ CD4 T cells and log-transfromed lung CFU at day 28 post-infection. The R^2^ value was calculated using a simple linear regression.

**Supplemental Fig 2. Vaccine-driven cytokine responses in the setting of pre-existing immunity.** (A) Parenchymal lung cells isolated from unvaccinated, CoMtb, CoMtb+H56/CAF01, or CoMtb+BCG-immunized mice were assayed for the production of different cytokine combinations upon ex vivo ESAT6 peptide stimulation 28 days post-Mtb challenge. (B-C) CB6F1 mice administered different Mtb-naive or post-Mtb exposure vaccines were quantified for cytokine production by parenchymal lung CD4 T cells following ex vivo ESAT6 peptide stimulation at day 98 post-Mtb aerosol challenge. Shown are (B) the percentage of parenchymal lung cells that co-produce IL-2 and IFNγ and (C) the quantification of IL-17-containing cytokine combinations.

**Supplemental Fig 3. Peripheral cytokine responses are augmented by Mtb exposure prior to aerosol challenge.** Mice were left unvaccinated, immunized with BCG or H56, or administered intradermal Mtb (CoMtb) followed by BCG or H56 immunization. Spleens were harvested 6 weeks following the final immunization for ex vivo peptide stimulation. (A,B) Total numbers of IFNγ+TNF+IL-2+ or IFNγ+TNF+ CD4 T cells in the spleens of mice following stimulation with (A) ESAT6 and (B) TB10.4 peptide, respectively (no IL-2 production was observed upon TB10.4 stimulation). (C) Representative flow cytometry plots of intracellular cytokine staining for IFNγ and TNF in the spleens of different vaccination groups following ex vivo ESAT6 peptide stimulation. (D-E) Spleens were also stimulated ex vivo with Ag85B(P25) peptide, and (D) the magnitude and quality of the response were measured by plotting the composition of cytokine-producing CD4 T cells that emerged. (E) The frequency of IL-2+ cells that co-express IFNγ was plotted.

**Supplemental Fig. 4. Lung antigen-specific CD4 T cells are expanded upon vaccination in the setting of pre-existing immunity.** (A) Proportion of parenchymal (IV-) CD4 T cells that produced IFNγ/TNF/IL-2 upon ESAT6 stimulation 6 weeks following vaccination. (B-C) Total numbers of IFNγ+TNF+IL-2+ CD4 T cells in the lung parenchyma of mice following stimulation with (B) ESAT6 or (C) TB10.4. (D) Representative flow cytometry plots of intracellular cytokine staining for IFNγ and TNF in the lungs of mice in different vaccination groups following ex vivo ESAT6 stimulation. (E-F) Representative flow cytometry plots of tetramer staining for (E) ESAT6-and (F) TB10.4-specific CD4 T cells 6 weeks following vaccination.

## Notes

### Competing Interest Statement

The authors have declared no competing interest.

