## Supplementary figures and images for "Subunit vaccination enhances protection conferred by prior *Mycobacterium tuberculosis* exposure"

### Supplemental Figures

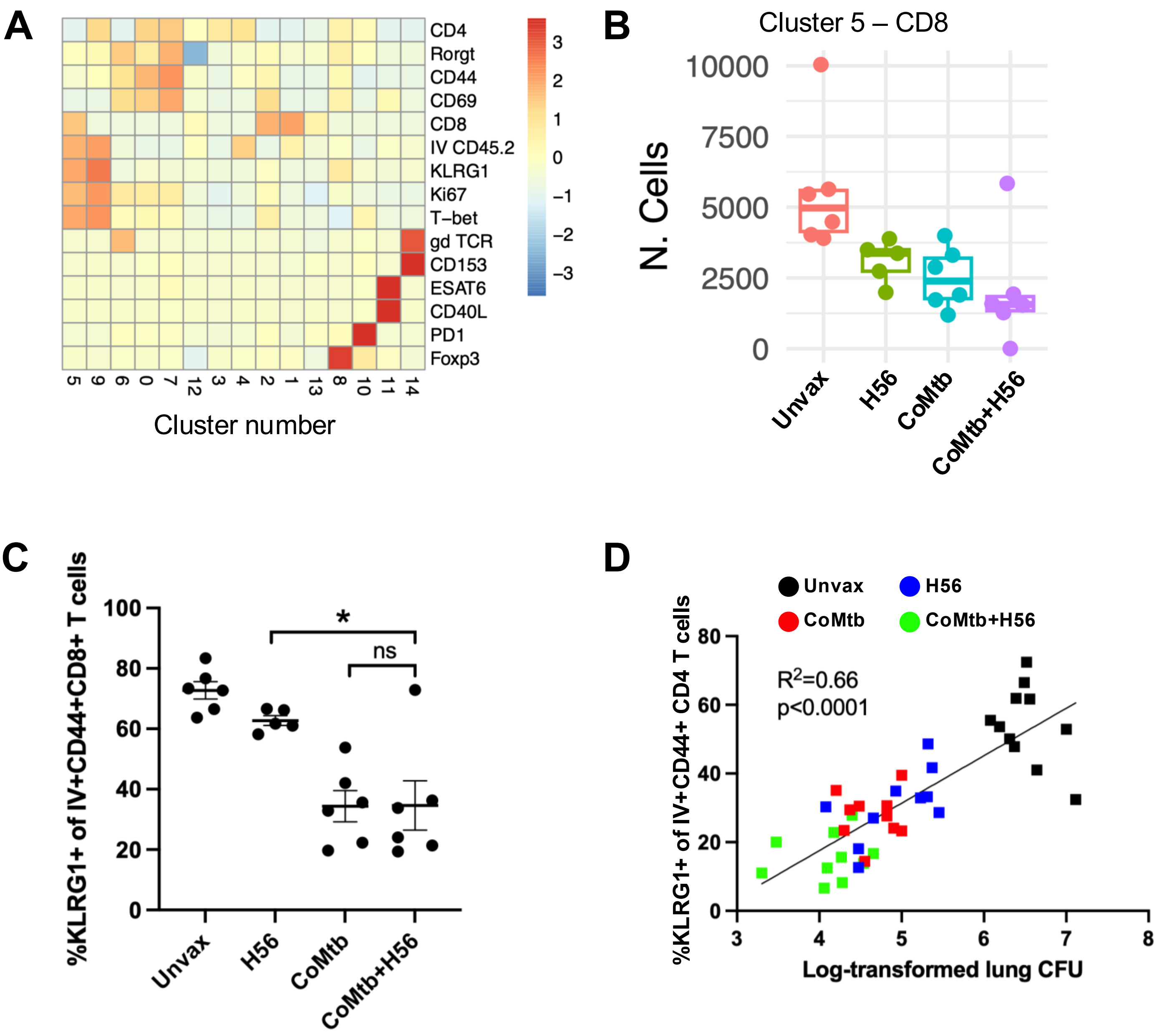

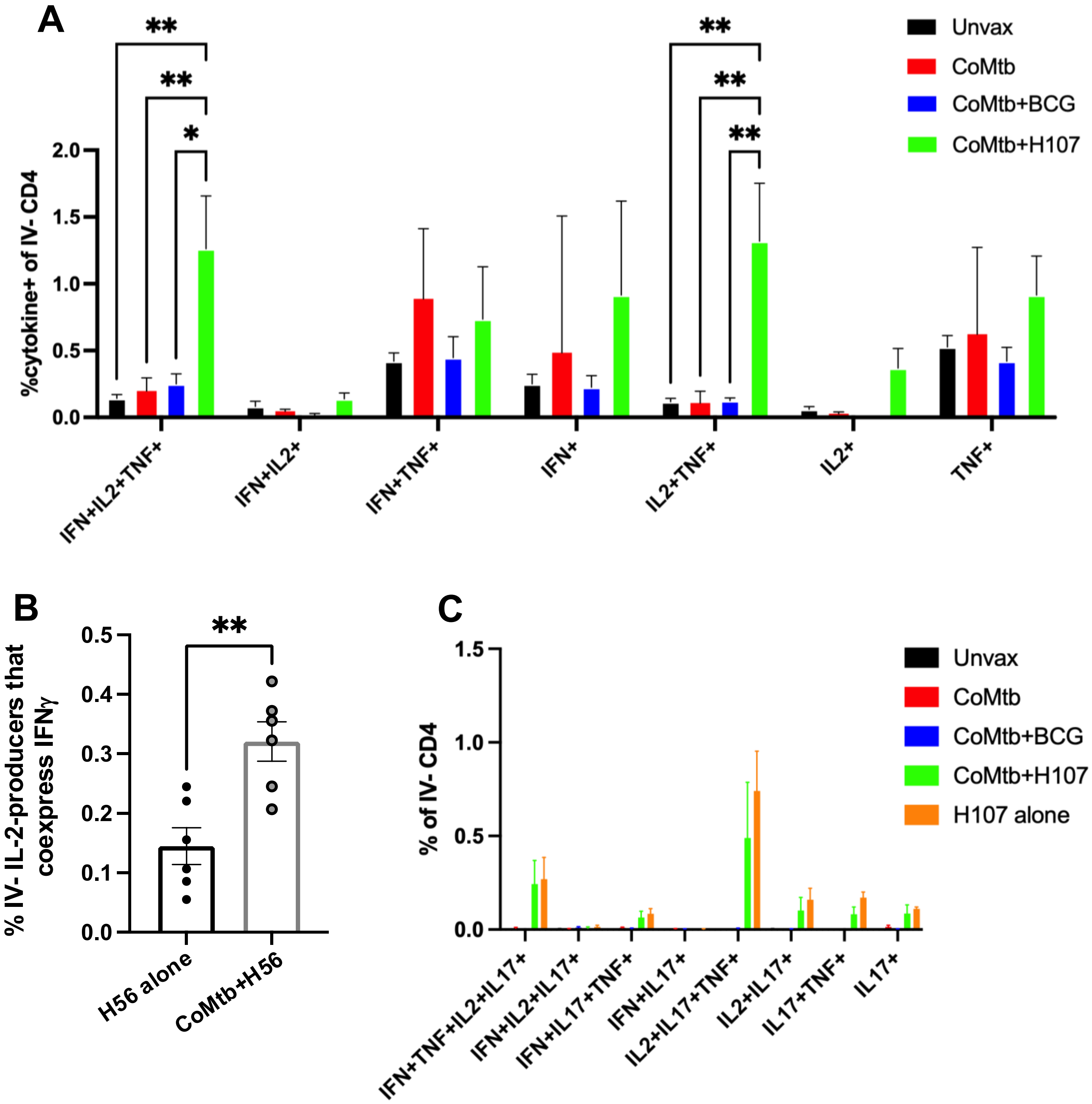

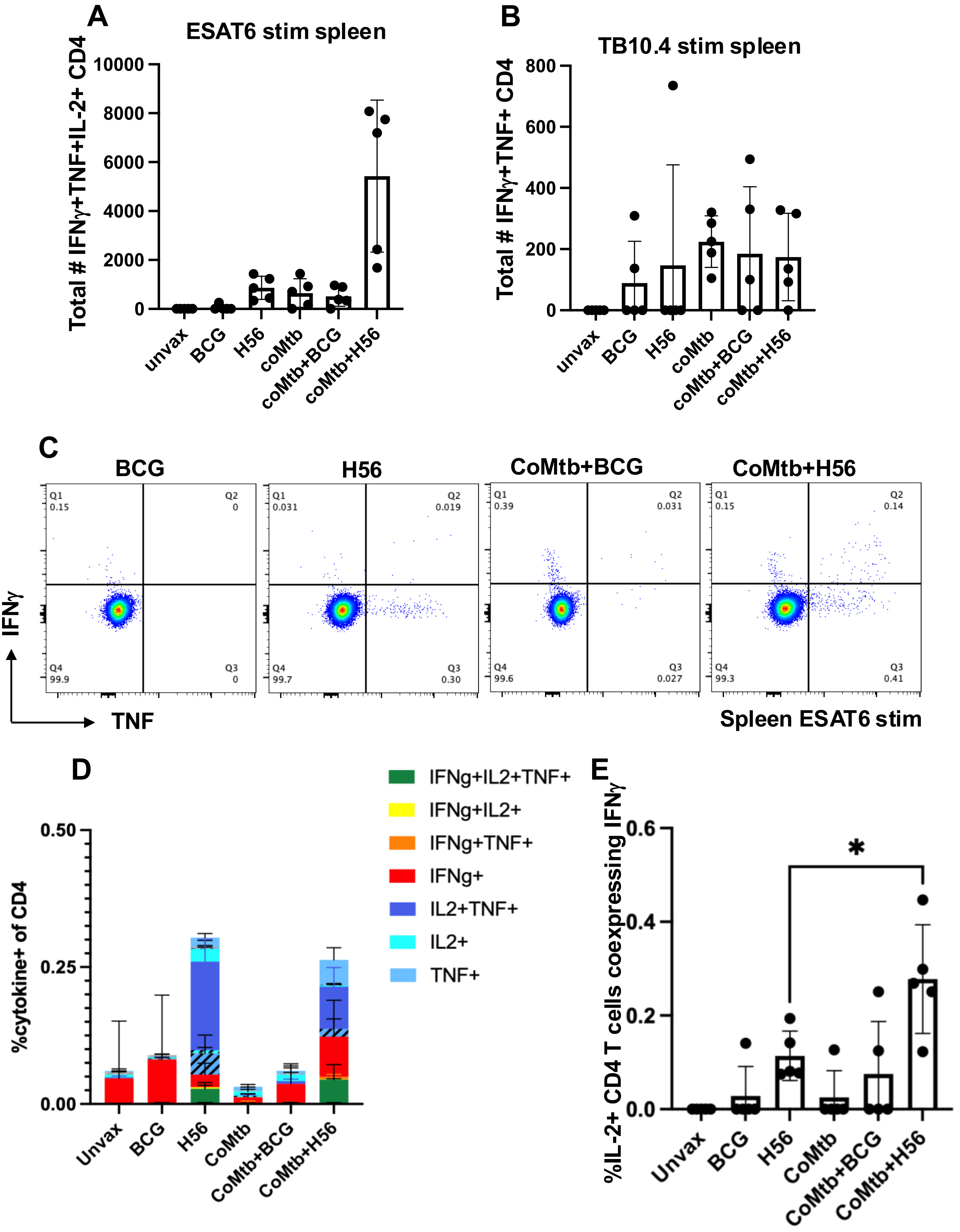

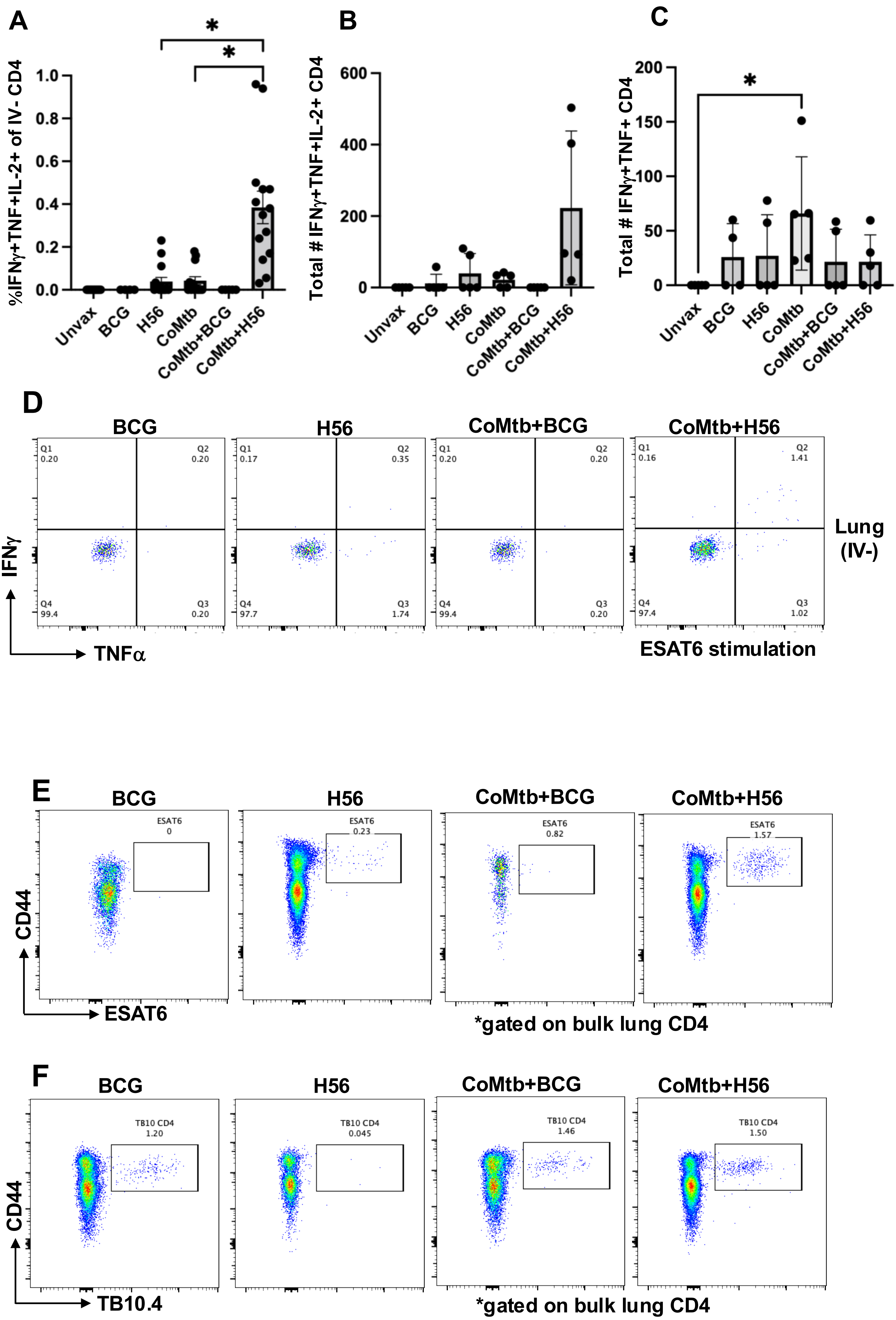
